# Anti-PTHrP blockade limits CD8+ T-cell exhaustion in anti-cancer immunotherapy

**DOI:** 10.1101/2024.10.23.619890

**Authors:** Bastien Moës, Yunfeng Gao, Ekaterina Demina, Richard Kremer, Christopher E. Rudd

## Abstract

Cancer is a major global health concern, with immune suppression hindering treatment. Immunotherapy, specifically immune checkpoint blockage on T cells, has revolutionized cancer treatment. T-cell exhaustion is an abnormal activation state that develops when continuous exposure to antigens, like cancer. In this context, recent evidence suggests that parathyroid hormone-related protein (PTHrP) plays a previously underappreciated role in fostering an immunosuppressive tumor microenvironment. Further, blocking PTHrP activity reduces primary tumor growth, prevents metastasis, and prolongs survival in mice with various cancers. Here, we confirm that administration of anti-PTHrP monoclonal antibodies can reduce the growth of B16-PDL1 melanoma tumors and that although the therapy did not alter the presence of CD4+ and CD8+ TILs, we noted that all stages of T-cell exhaustion were reduced. Further, the expression of cytolytic proteins PERFORIN and GZMB also increased. By contrast, anti-PTHrP therapy increased the relative presence of pre-pro B cells with a decline in mature B cells in the bone marrow. Overall, our data indicates that anti-PTHrP therapy acts by reducing T-cell exhaustion and by affecting B-cell development. These provide further mechanistic evidence to support the application of anti-PTHrP blockade as an alternate therapeutic approach to boost anti-tumor immunity.

## Introduction

Cancer remains one of the leading causes of mortality worldwide, with tumor-induced immune suppression posing a significant obstacle to successful treatment^1^. Immunotherapy has revolutionized cancer treatment by harnessing the body’s immune system to fight malignant cells^2^. Being a promising strategy that targets T cell exhaustion, immune checkpoint blockage (ICB) of negative co-receptors on T cells has therefore emerged as a key cancer treatment objective. In this context, the blockade of PD-1 and PD-L1 with monoclonal antibodies in immune checkpoint blockade (ICB) has shown therapeutic benefits in patients with various cancers. These tumors include non-small cell lung carcinoma (NSCLC), melanoma, and bladder cancer^3-6,4^. Response rates have been observed with anti-PD-1 alone, or in combination therapy with anti-cytotoxic T-lymphocyte–associated antigen 4 (CTLA-4) ^4,7-9^. LAG-3 is another “checkpoint” that cancer can exploit. Drugs like relatlimab are being tested to block LAG-3 in combination with PD-1 inhibitors. Not all patients, meanwhile, benefit from these therapies, which emphasizes the necessity for additional or substitute immunotherapy strategies^10^.

One key issue related to the success of ICB is T cell exhaustion that results from continuous exposure to antigens, like cancer. This dysfunctional statement results in epigenetic and transcriptional changes that impair CD8+ T cell activation. Through the overexpression of inhibitory receptors like PD-1 and CTLA-4, decreased cytokine production and their proliferation, and an inability to effectively destroy target cells, exhausted T cells lose their capacity to mount an effective immune response against cancer cells^10-12^. In certain models, stem-like (Tpex) CD8+ T cells (PD1+TCF-1+TIM3-) develop into terminally exhausted T cells (PD1+TCF-1-TIM3+)^13,14^.

Despite the success of ICB, as mentioned, a significant proportion of patients are not cured, emphasizing the ongoing need for a better understanding of checkpoint blockade mechanisms to facilitate the development of more effective therapies. Recent evidence suggests that parathyroid hormone-related protein (PTHrP), a protein commonly associated with hypercalcemia of malignancy is frequently overexpressed in various cancers, including breast, lung, and prostate cancers, where it has been shown to promote tumor progression through paracrine and autocrine mechanisms^15,16^. Recent data demonstrate that blocking PTHrP activity reduces primary tumor growth, prevents metastasis, and prolongs survival in mice with breast cancer, melanoma pancreatic ductal adenocarcinoma cancer (PDAC), one of the most aggressive forms of cancer^17-19^. PTHrP has been linked to a variety of functions, including intracellular calcium, proliferation/hypertrophy, and differentiation. These functions are most likely the result of the mature PTHrP protein being processed into at least three active peptides, each of which is assumed to act independently via autocrine, paracrine, and intracrine pathways^20,21^. PTHrP isoforms are produced and secreted by malignant cells, including moieties with intact N-termini as well as mid-region fragments. The most widely studied N-terminal peptide, PTHrP(1–34), binds to the parathyroid hormone receptor 1 (PTHR1). PTHrP(1-34)-PTHR1 binding starts signaling cascades (e.g., extracellular signal-regulated kinase (ERK), AKT, cyclin D1, RUNX, and cAMP) that ensue in the direction of cell survival and proliferation^22-25^. Crucially, by a process connected to LAMP1 signaling, PTHR1 ablation in osteoprogenitor from bone marrow hinders proper B cell development and affects mature B cell retention in bone marrow ^**26**^.

Considering these findings, targeting PTHrP represents a novel therapeutic avenue for boosting T and B cell function and reducing tumor growth. Our data provides evidence that anti-PTHrP therapy acts by slowing the progression of CD8+ tumor-infiltrating lymphocytes (TILs) to a non-functional exhausted state and influencing B-cell development. This further elucidates the mechanism and supports exploring anti-PTHrP blockade as an alternative therapeutic approach to enhance anti-tumor immunity.

## Results

### Anti-PTHrP therapy limits tumour growth and T cell exhaustion

To examine the impact of anti-PTHrP therapy on tumor control in another tumor model (melanoma) and the phenotype of tumor infiltrating lymphocytes (TILs), mice were implanted with B16DL1 cells, and 200 ug of anti-PTHrP antibody was injected every 2 days starting on the 5th day following the transplantation (Fig. 1A). As showed by lower volume, tumor control under anti-PTHrP treatment was demonstrated to be more effective than in the control group (Ctrl) (Fig. 1B). When focusing on TILs, we observed no obvious effect of anti-PTHrP therapy on CD8+ or CD4+ T cell infiltration, according to TIL extracted from B16PDL1-implanted mice, as no difference in their relative proportion was observed when compared with Ctrl mice (Fig. 1C). Exhausted CD8+ T cell presence in TIL is known to be correlated with tumor growth^27^. We thus investigated exhaustion lineage. This analysis revealed first that the PD-1+ population among CD8+CD44+ T cells was greatly decreased under anti-PTHrP treatment. Concomitantly, stem-like (PD-1+TCF-1+TIM3-) and exhausted (PD1+TCF-1-TIM3+) cells belonging to the CD8+ exhausted population were found to be significantly reduced when B16PDL1-implanted mice were treated with an anti-PTHrP antibody (Fig. 1D). Apart from the decline in T cell exhaustion, we also observed that the anti-PTHrP treatment lowered the expression of two inhibitory receptors, PD-1 and TIM3, and TOX, the master transcription factor of exhaustion lineage establishment (Fig. 1E and Fig. 1F). Lastly, it was observed that anti-PTHrP therapy significantly increased the production of two cytotoxic proteins involved in the death of tumor cells, PERFORIN (PRF) and GRANZYM B (GZM B), in CD44+CD8+ TIL (Fig. 1G). All things considered, these findings show that anti-PTHrP therapy reduced tumor growth (at least in part) by limiting T cell exhaustion and super-armed CD8+ TIL and suggest anti-PTHrP as a viable immunotherapy treatment option.

**Figure 1:**
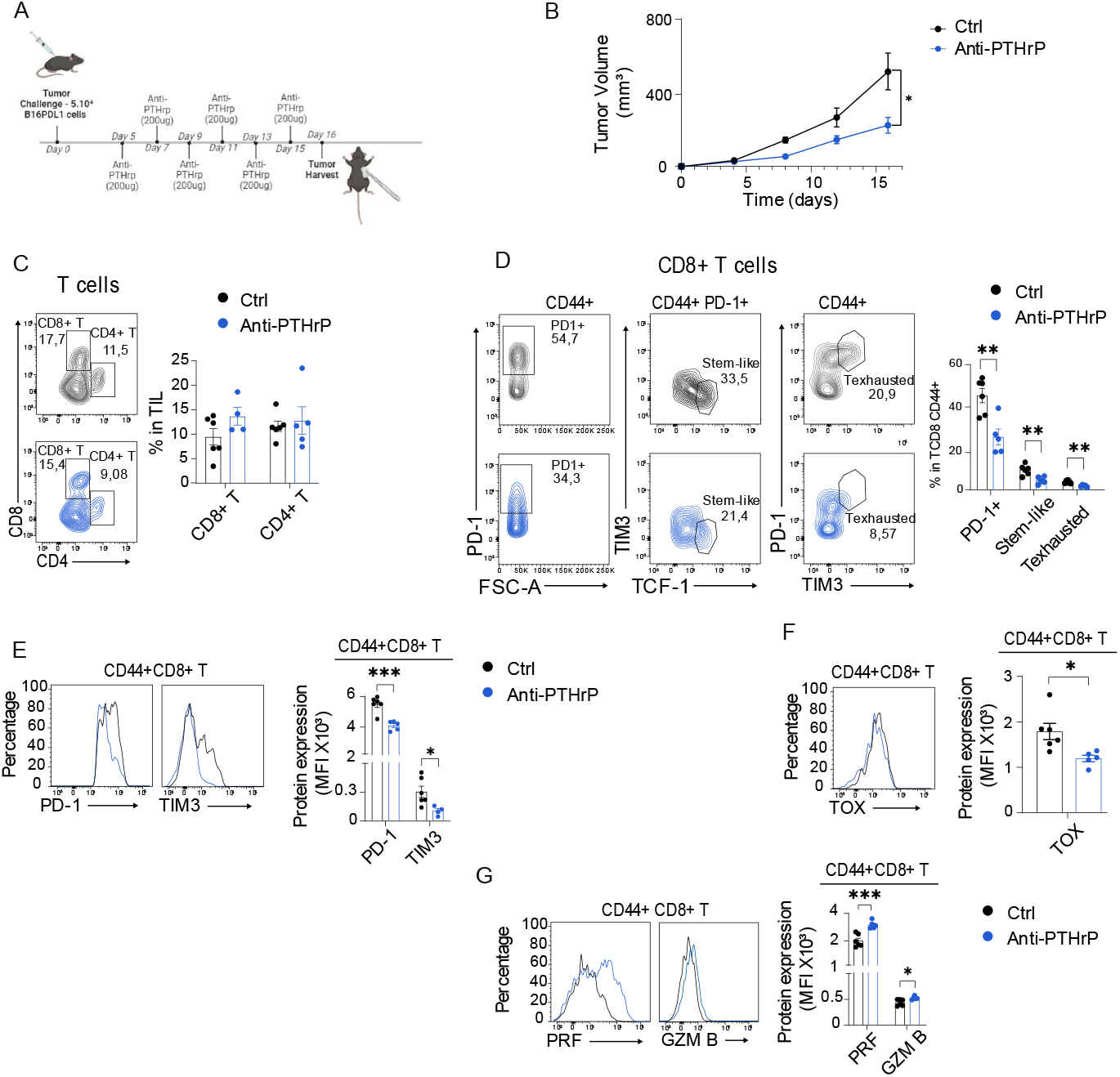
Anti-PTHrP therapy limits tumour growth and T cell exhaustion. **A**. Experimental layout: mice were implanted with 5.10^4^ B16PDL1 cells and injected with 200 ug of anti-PTHrp antibody every 2 day from day 5 post-implantation. **B**. Quantification of B16PDL1 tumor volume without or with anti-PTHrP injection; n=5. Data are representatitve from 3 independent experiments **C**.Representative flow cytometry plots and quantification of proportion of CD4+ and CD8+ TIL : n=5. Data are representatitve from 3 independent experiments **D**. Representative flow cytometry plots and quantification of proportion of PD1+, Stem-like (PD1+TCF-1+TIM3-) and Texhausted (PD1+TCF-1-TIM3+) population among CD44+ CD8+ TIL: n=5. Data are representatitve from 3 independent experiments **E**. Representative flow cytometry plots and quantification of PERFORIN and GRANZYME B (GZM B) expression in CD44+CD8+ TIL. **F**. Representative flow cytometry plots and quantification of TOX, PD-1 and TIM3 expression in CD44+CD8+ TIL: n=5. Data are representatitve from 3 independent experiments Data are shown as mean ± SEM. Statistics are analyzed by unpaired t test; ∗p < 0.05, ∗∗p < 0.01, ∗∗∗p < 0.001, and ∗∗∗∗p < 0.0001. NS, not significant.

### Anti-PTHrP influences B cell development and mature B cells presence in bone marrow

PTHrP binds to PTH1R, a receptor found to play an important role during B cell development in bone marrow^26^. Moreover, B cells have been strongly characterized as an CD8+ TIL activator and are important for tumor cell elimination^28,29^. We next explored B cell development in bone marrow using Hardy’s fraction nomenclature (Fig. 2A). Tumor-implanted mice treated with anti-PTHrP showed an interesting difference during B cell development. First, the presence of cells from Fr.A was found to be significantly increased following anti-PTHrP treatment and is consistent with previous reports^26^. Impressively, mice treated with anti-PTHrP antibody showed a great reduction of cells from Fr.F, also known as mature B cells (Fig. 2B). Of note, mature B cells are not produced in bone marrow but are instead coming from the spleen and blood. All of these findings indicate that anti-PTHrP therapy influences the B cell lineage as well. They also suggest that under anti-PTHrp therapy, mature B cells may be stopped in a different lymphoid organ where they may have a crucial involvement in tumor regression.

**Figure 2:**
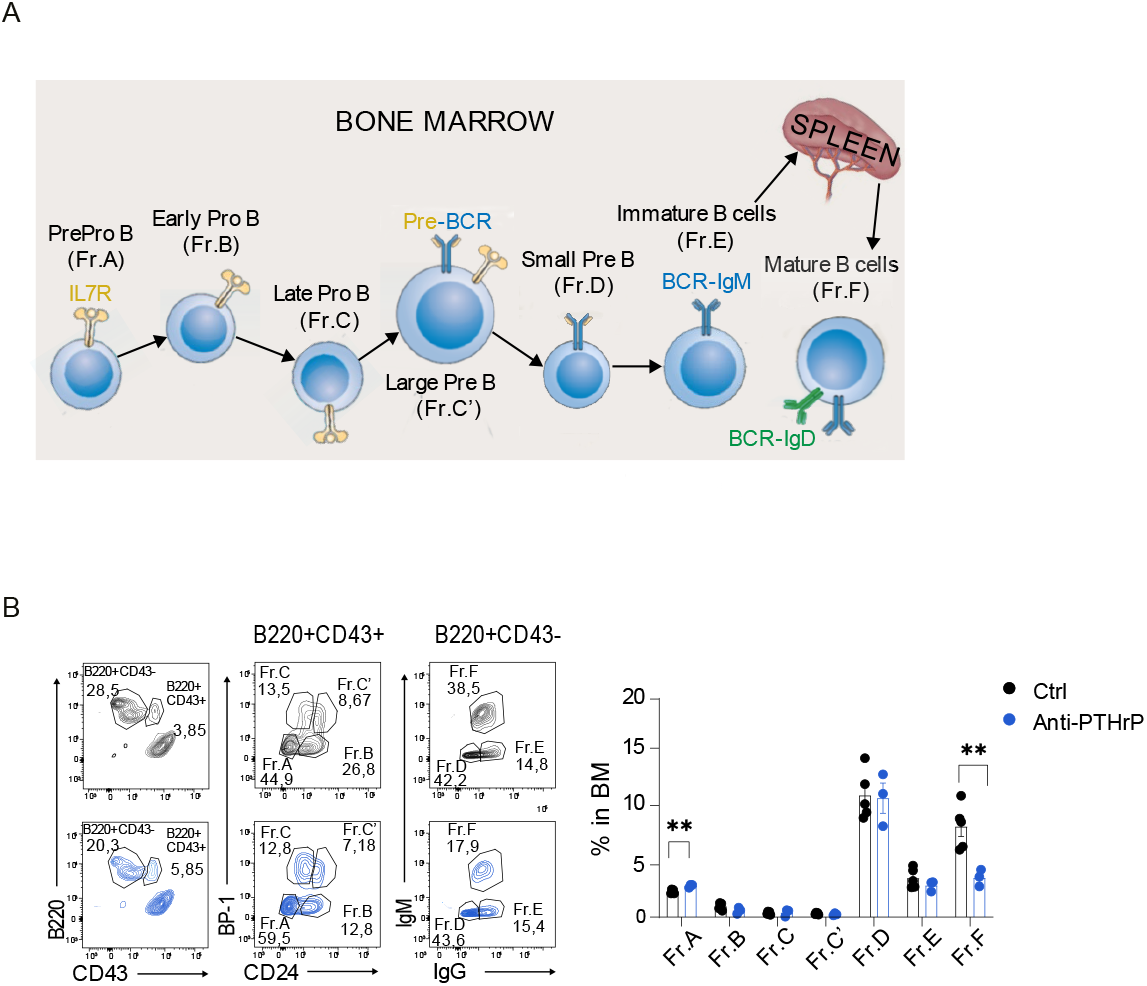
PTHrP blockade influences B cell development and mature B cells presence in bone marrow. **A**.Representation of Hardy’s fraction in bone marrow : nomenclature. **B**. Representative flow cytometry plots and quantification of proportion of B cell fraction (Fr.A-F) in bone marrow: n= 4-3. Data are shown as mean ± SEM. Statistics are analyzed by unpaired t test; ∗p < 0.05, ∗∗p < 0.01, ∗∗∗p < 0.001, and ∗∗∗∗p < 0.0001. NS, not significant.

## Discussion

PTHrP was earlier shown to be a great, underappreciated targetable protein that could inhibit tumors even in PDAC, one of the most aggressive types of cancer^17-19^. In this context, PTHrP has been shown to be a driver of disease development; inhibiting PTHrP activity in mice decreases the growth of primary tumors, stops metastasis, and increases their survival time. Nevertheless, to the best of our knowledge, there has never been any evidence linking PTHrP to immune cell activation. Here, using the B16 model of melanoma cancer, we provide evidence that anti-PTHrP blockade acts by slowing the progression of CD8+ tumor-infiltrating lymphocytes (TILs) to a non-functional exhausted state and by influencing B-cell development. This further elucidates the mechanism and supports exploring anti-PTHrP blockade as an alternative therapeutic approach to enhance anti-tumor immunity.

Initially, we showed that anti-PTHrP was effective in the B16-F10 melanoma model expressing PDL1. Surprisingly, unlike other forms of checkpoint blockade such as anti-PD-1, anti-PTHrP failed to promote an increased presence of CD4 and CD8+ TILs in the tumor. This suggests that anti-PTHrP has no effect on the migration of T-cells into tumors or in the replication of cells once in the tumor microenvironment.

Nevertheless, despite this, there was a striking effect on the exhaustion phenotype of the CD8+ TILs. inhibition was also found to restrict T cell exhaustion lineage establishment by lowering both stem-like and exhausted T cell presence in tumors. Additionally, we demonstrated that PTHrP blockade not only modulates the expression of inhibitory receptors (such as PD-1 and TIM3) on CD8+ TILs but also acts as a mediator of the exhaustion lineage by controlling TOX, the master TF of the exhaustion lineage^30^. In keeping with the limiting effect on exhaustion this, we also found that anti-PTHrP therapy increased the expression of GZM B and PRF, two proteins involved in the process of killing tumor cells^31,32^. These findings provide the first proof that PTHrP inhibition can effectively regulate tumor development by boosting CD8+ TIL activation.

Furthermore, we showed that anti-PTHrP therapy also influenced the presence of B-cells. Previous studies reported that ablation of PTHR1 alters B cell development in bone marrow^26^. In tumor context, we observed that blocking PTHrP activity also impairs early B cell development. Unlike what was previously reported, we discovered that mature B cell retention in bone marrow is decreased rather than enhanced following anti-PTHrP treatment. Two possible explanations for this might be that in the previous study ablation of PTHR1 was inherent to osteoprogenitor from bone marrow or that parathyroid hormone (PTH) is another important substrate of PTHR1, and so the signaling through this receptor is still occurring. We hypothesize that the decreased abundance of mature B cells in bone marrow during anti-PTHrP treatment could be due to their sequestration in distinct lymphoid organs, where they may contribute to tumor regression through mechanisms that have yet to be fully elucidated. This observation opens intriguing possibilities regarding the involvement of B cells in the context of PTHrP inhibition and highlights the need for further research to elucidate their roles during immunotherapy.

In summary, this study provides new compelling innovative evidence that anti-PTHrP antibodies reduce, at least in part, tumor growth by reversing T cell exhaustion and strongly suggests that PTHrP inhibition represents a promising therapeutic strategy for enhancing anti-tumor immunity. Additionally, our findings raise the possibility that B cells may have a previously unrecognized role in this process, particularly under anti-PTHrP therapy, potentially contributing to tumor regression through mechanisms occurring in secondary lymphoid organs. These insights offer new avenues for understanding the immunological effects of anti-PTHrP therapy and suggest novel therapeutic combinations that target both T and B cells in cancer treatment. This is particularly relevant considering the multifaceted roles of B cells in cancer, such as CD8+ TIL activation and tumor cell antibody production^33^.

## Methods

### Sex as a biological variable

In this study, both male and female mice were used in the animal model experiments to investigate the effects of anti-PTHrP therapy on B16-PDL1 tumor regression. However, sex was not considered as a biological variable in any of the analyses. *Mice*. 8 weeks to 12 weeks old C57BL/6J (Strain #:000664) mice from the Jackson Laboratory were used in this study. All mice were housed in an animal facility with a 12 h light/12 h dark cycle and had free access to food and water throughout the study period. All mouse studies were authorized by the Animal Care Use and Review Committee of CR-HMR.

### B16-PDL1 implantation, anti-PTHrP antibody injection and TILs isolation

B16-PDL1 tumor cells were cultivated in DMEM + 10% until reaching 60% to 90% of confluence and then subcutaneously implanted in mice. Anti-PTHrP antibody (IgG3; 1-33) from Dr Richard Kremer’s lab (McGill university) were diluted at 1mg/ml and 200ug were IP injected in mice each 2 days from d5 post-implantation until tumor extraction. Tumors were smashed on 70 um strainers in RPMI on ice. Tumor lysate was spined down at 350g for 5min and resuspended in PBS. PBS containing tumor lysate was add to lymphocytes separation medium (Corning #07421010) and TILs were separated and isolated after centrifugation at 2000 rpm for 20 min at RT. TIL were stained as described in the specific section.

### Cell isolation

Bone marrow was taken from femurs and tibias of 8- to 16-week-old mice. Tibias and femurs were first flushed with PBS/2mM EDTA + 2% FBS, filtrated on a 70µm membrane and centrifuged 5min at 350g. Then, pellet was resuspended in 5 ml of red blood lysis buffer (Biolegend #420302) 5 min at RT. Lysis reaction was stopped by adding 10ml of PBS/2mM EDTA + 2% FBS and the resuspended solution was centrifuged 5min at 350g.Spleen was taken from 8- to 16-week-old mice and pressed in a 6 wells plate containing 2ml of red blood lysis buffer (Biolegend #420302) 2 more ml were added and then incubated 5min at RT. Lysis was stopped with 20ml of PBS/2mM EDTA and the solution was filtered on a 70µm membrane and then centrifuged 5min at 350g. Pellets from spleen cells were resuspended in 1ml of PBS/2mM EDTA + 2% FBS. Alive cells from bone marrow and spleen were counted on a Neubauer counting chamber (Hirschmann #8100103) with Trypan blue (Thermoscientific # SV30084.01).

### Facs staining and immunophenotyping

Isolated cells from spleen and bone marrow were stained with Fixable Viability Dye eFluor™ 780 (dilution:1/1000; ebioscience # 65-0865-18) 20min at 4°C and then blocked with 100µl of PBS/2mM EDTA + FBS 5% + 2,5 ng/ml purified rat anti-mouse CD16/CD32 (BD Biosciences #553141) 20min at 4°C. Cells were then centrifuged 5min at 350g and stained with primary antibodies for 30min at 4°C. The fluorochrome-conjugated antibodies were as follows for flow cytometry: BV605 Rat Anti-Mouse CD4 (RM4-5/RM4-4; Biolegend #116027), PE-Cy™7 Rat Anti-Mouse CD8a (53-6.7; BD Biosciences #552877), Alexa Fluor® 700 Hamster anti-Mouse CD3e (500A2; BD Biosciences #557984), PerCP-Cy™5.5 Rat Anti-Mouse/human CD44 (IM7; Biolegend #103032), BD OptiBuild™ BV605 Mouse Anti-Mouse CD366 (TIM-3) (5D12; BD Biosciences #7476241), Brilliant Violet 510™ anti-mouse CD279 (PD-1) Antibody (29F.1A12; Invitrogen; 35-5893-80, Alexa Fluor® 700 anti-mouse CD90.2 (Thy-1.2) Antibody (30-H12; Invitrogen #25-1011-80) PerCP/Cyanine5.5 anti-mouse IgM Antibody (RMM-1; Biolegend #406512), CD24 Monoclonal Antibody, Brilliant Ultra Violet™ 737 (M1/69; ebioscience #367-0242-80), BV421 Rat Anti-Mouse CD249 (F344; Biolegend #740013), APC anti-mouse CD43 Antibody (S11; Biolegend #143208), Brilliant Violet 711™ anti-mouse/human CD45R/B220 Antibody (RA3-6B2; Biolegend #103255), FITC anti-mouse IgD Antibody (11-26c.2a; Biolegend #405704). Cells were fixed with 2% PAF (Corning #14-959-1A) 15min at 4°C, washed 2 times in PBS/2mM EDTA +FBS 2% and then nuclear permeabilized with FOXP3 kit (Thermofisher #00-5523-00) for 30 min at 4°C. Cells were stained with antibodies targeting intracellular protein for 50min at RT and diluted in PBS/2mM EDTA + FBS 5%. The fluorochrome-conjugated antibodies were as follows for flow cytometry: Alexa Fluor® 647 ConjugateTCF1/TCF7 Rabbit mAb (C63D9; Cell signaling # 6709), PE anti-human/mouse Granzyme B Recombinant Antibody (QA16A02; Biolegend # 135008), APC anti-mouse Perforin Antibody (S16009B; Biolegend # 154404), TOX Monoclonal Antibody PE (TXRX10; ebiosciences # 12-6502-82). Cells were washed 2 times in PBS/2mM EDTA +FBS 2%. BD FacsFortessa was used for cell phenotyping and intracellular detection. Mean fluorescence intensity (MFI) of specific markers was quantified on T cell subpopulations using FlowJo.

### Statistics

GraphPad Prism version 8 (GraphPad Software) and Flowjo (Tree Star, Ashland, USA) were used for all statistical analyses. The sample size for the experiments was 3 or more. Results are represented as the MFI ± SEM unless otherwise indicated. Comparisons between 2 groups were conducted using an unpaired t test. All reported P values are 2 sided and were considered statistically significant at a P value of less than 0.05: ∗p < 0.05, ∗∗p < 0.01, ∗∗∗p < 0.001, and ∗∗∗∗p < 0.0001. NS, not significant.

## Author contributions

BM and CER conceived the project. BM, YG, ED, and CER designed the study. BM, YG and ED were involved in data acquisition. BM performed the statistical analysis. BM, RK and CER were involved in the interpretation of data. BM, YG and ED were responsible for the mouse colonies. RK provided anti-PTHrp antibody. BM, KR, and CER prepared the manuscript.

## Acknowledgements

We thank Facs operator from the Flow Cytometry Core Facility of CR-HMR for their help. CER was supported by the Canadian Institutes of Health Research Foundation grant (159912) (PI) and National Institutes of Health grant RO1 AI049466 (co-PI). RK was supported by supported by the Canadian Institutes of Health Research (MOP-142287) and is the inventor of the therapeutic monoclonal anti-PTHrP antibodies used in this study and authorized for use by Biochrom Pharma. RK holds an equity stake in Biochrom Pharma.Facs analysis were performed thanks to the Flow Cytometry Core Facility of CR-HMR.

Schematic illustrations were created using BioRender.com.

## Footnotes

All other authors declare no potential conflicts of interest.

